## Supplementary Figures for "Ongoing shuffling of protein fragments diversifies core viral functions linked to interactions with bacterial hosts"

### Supplementary Tables & Figures

**Table S1.** List of original PHROG annotations and the alternative names.

**Table S2.** List of odds ratios and p-values for each ECOD domain tested for being over-represented in mosaic rHMMs (Figure 2C and 2D).

**Table S3.** List of odds ratios and p-values for each functional class tested for being over-represented in mosaic rHMMs (Figure 4).

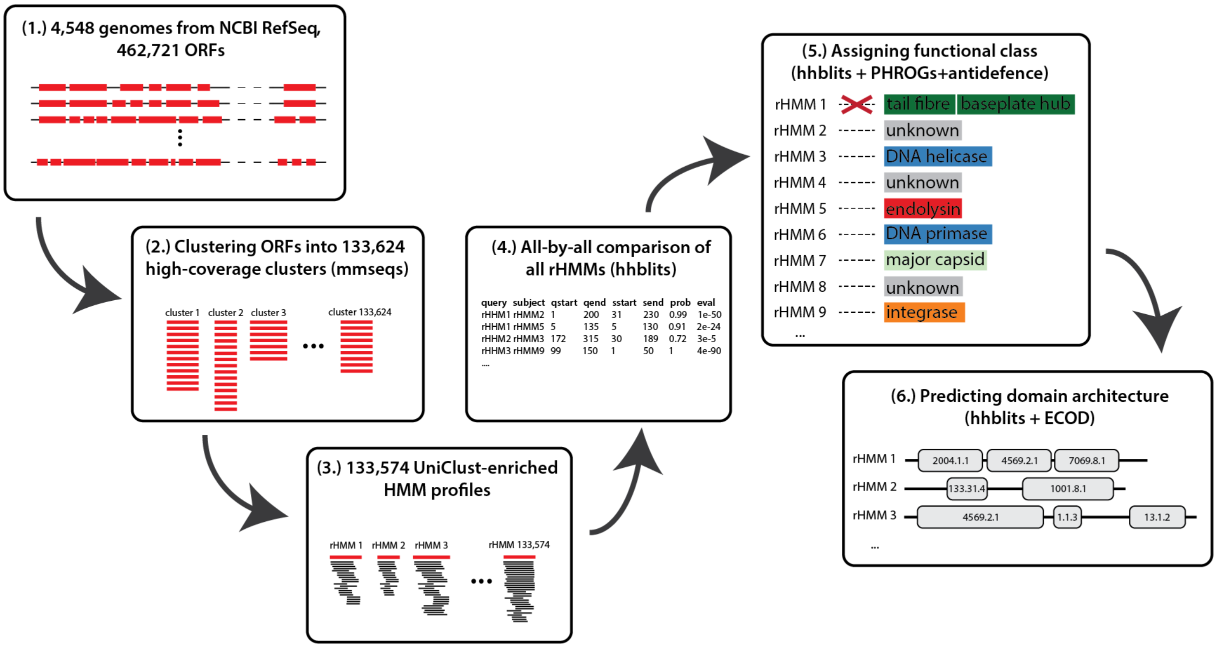


**Figure S1. Summary of methodology.** (1) We predicted 462,721 protein sequences in all phage genomes downloaded from NCBI RefSeq. (2) These protein sequences were clustered into 133,624 high-coverage clusters with `mmseqs2`. (3) A representative sequence from each cluster (with no more than 10 unknown characters) was taken and we aligned the UniClust30 database against it using `hhblits` to create 133,574 rHMMs (i.e., HMM profiles of representative protein sequences). All rHMMs were then compared against (4) each other using `hhblits` to generate the all-by-all comparison table, (5) PHROGs HMM database to assign a functional class to each rHMM, enhanced by hits to a database of anti-defence functions (Samuel & Burstein 2023), with conflicting assignments discarded, and (6) ECOD HMM database to predict protein domain architectures in each rHMM.

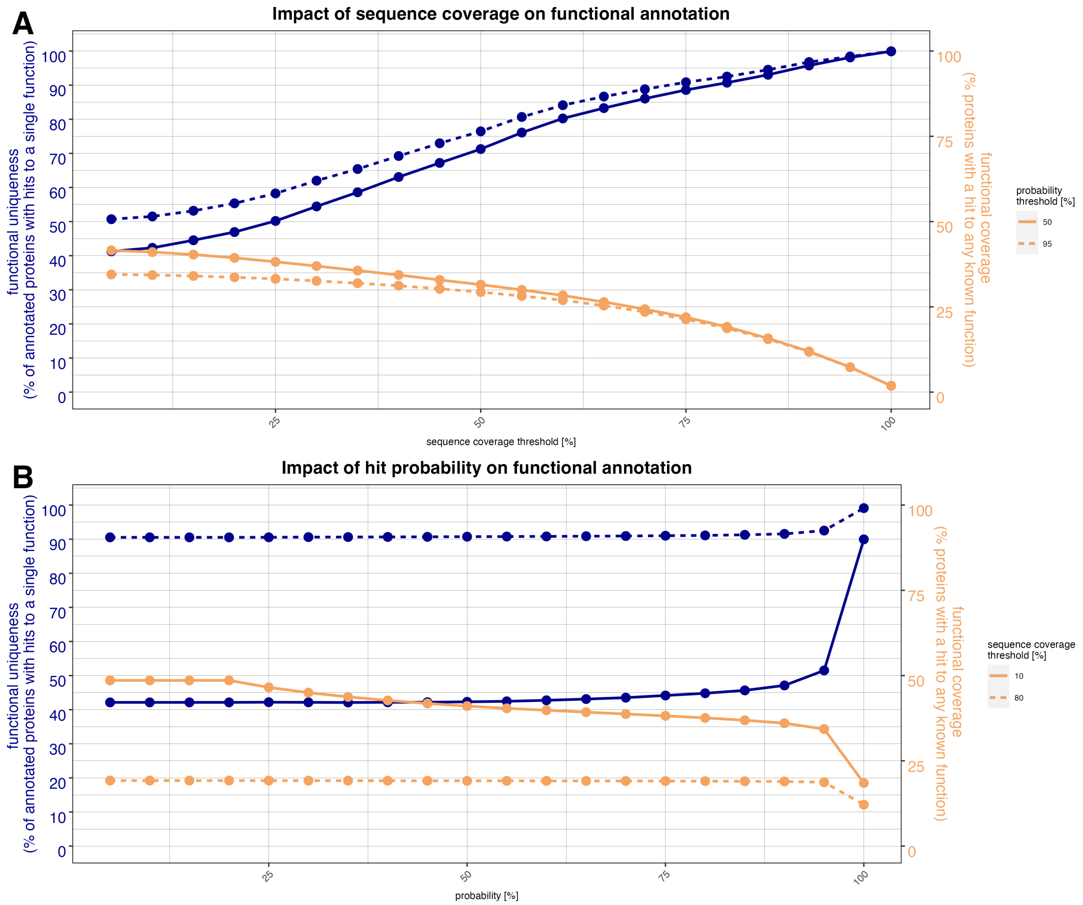


**Figure S2. Ambiguity of functional annotation increases with decreasing coverage.** Impact of `hhblits` parameters on functional annotation. (A) Functional uniqueness (percentage of annotated rHMMs with hits to a single function; blue) and functional coverage (percentage of rHMMs with hits to any known function; orange) as a function of sequence (both query and subject) coverage threshold, for different values of the probability hit threshold (solid vs. dashed line). (B) Functional uniqueness and functional coverage as a function of probability threshold, for two extreme values of sequence coverage threshold (both query and subject; solid vs. dashed line).

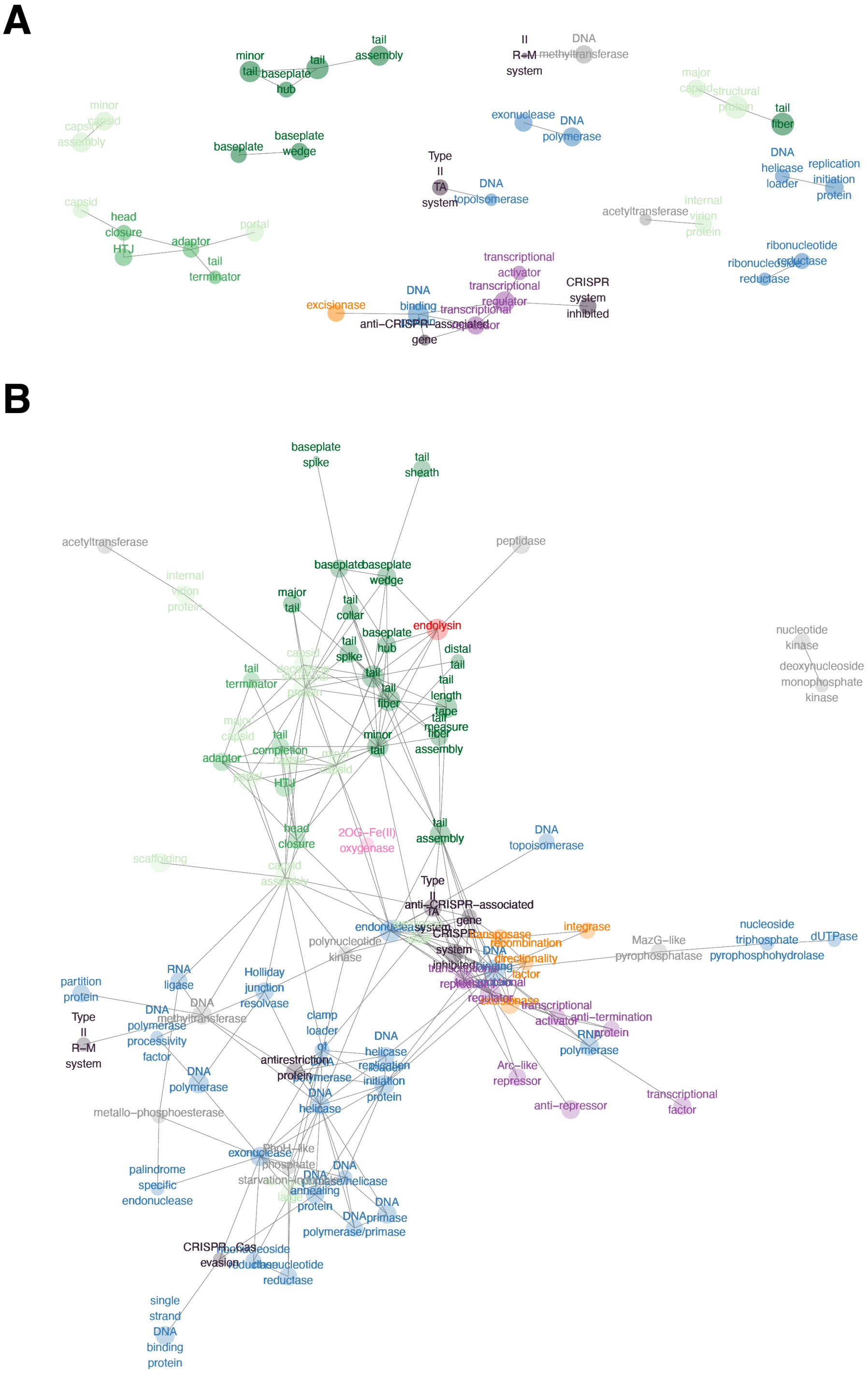


**Figure S3. Functional co-occurrence at a high and low sequence coverage threshold.** Each node represents a PHROG functional class. Nodes are linked an edge if they co-occurred at 80% (A) or 30% (B) coverage in at least 20 rHMMs at minimum 95% probability thresholds in hhblits. Colours refer to PHROG categories and node sizes refer to the number of rHMMs annotated by each PHROG functional class.

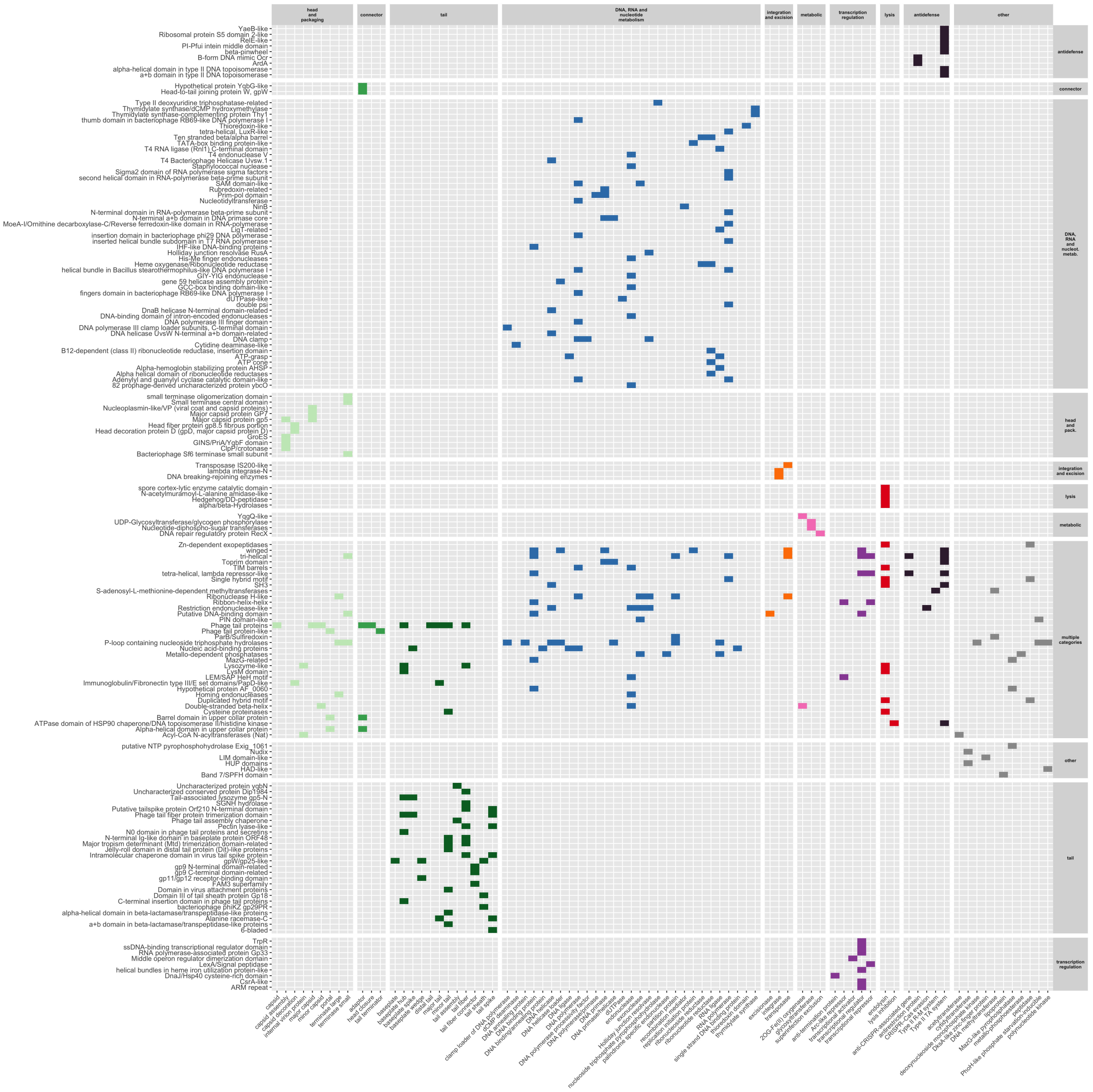

**Figure S4. Occurrence of different topologies in common functional classes.** Heatmap showing which folds (Y-axis), defined by ECOD domains (T-groups), are found in which common PHROG classes (X-axis). Colours refer to PHROG categories.

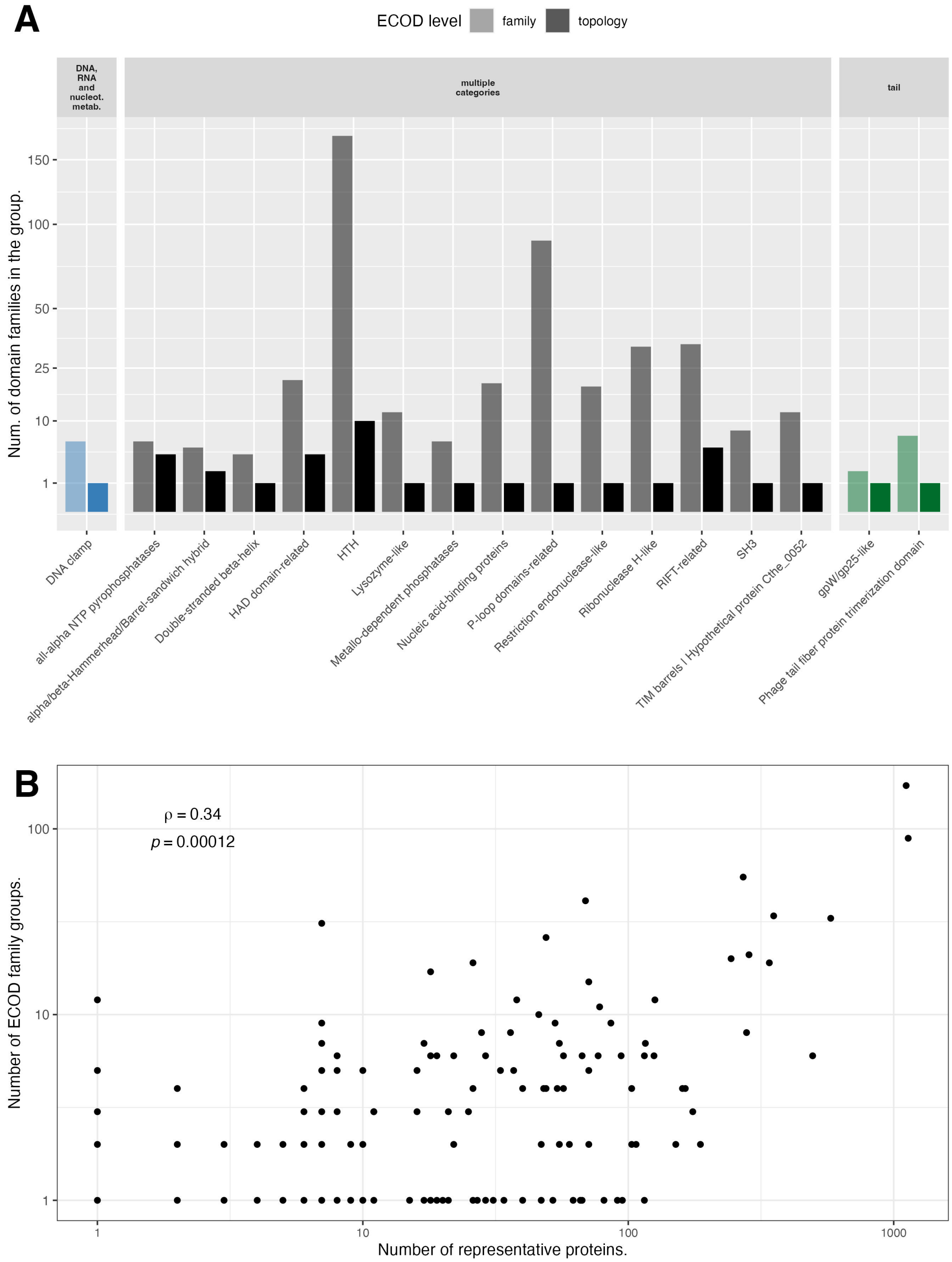


**Figure S5. Diversity and frequency of domains.** (A) Number of distinct domain families included in each H-group and detected within our rHMMs with valid functional annotations. Only the groups shown in Figure 1 are considered; (B) Number of distinct domain families included in each H-group and detected within rHMMs with known annotations vs. the number of rHMMs where a domain from given H-group is detected. Each point is an H-group present in our data, including the groups not shown in Figure 1.

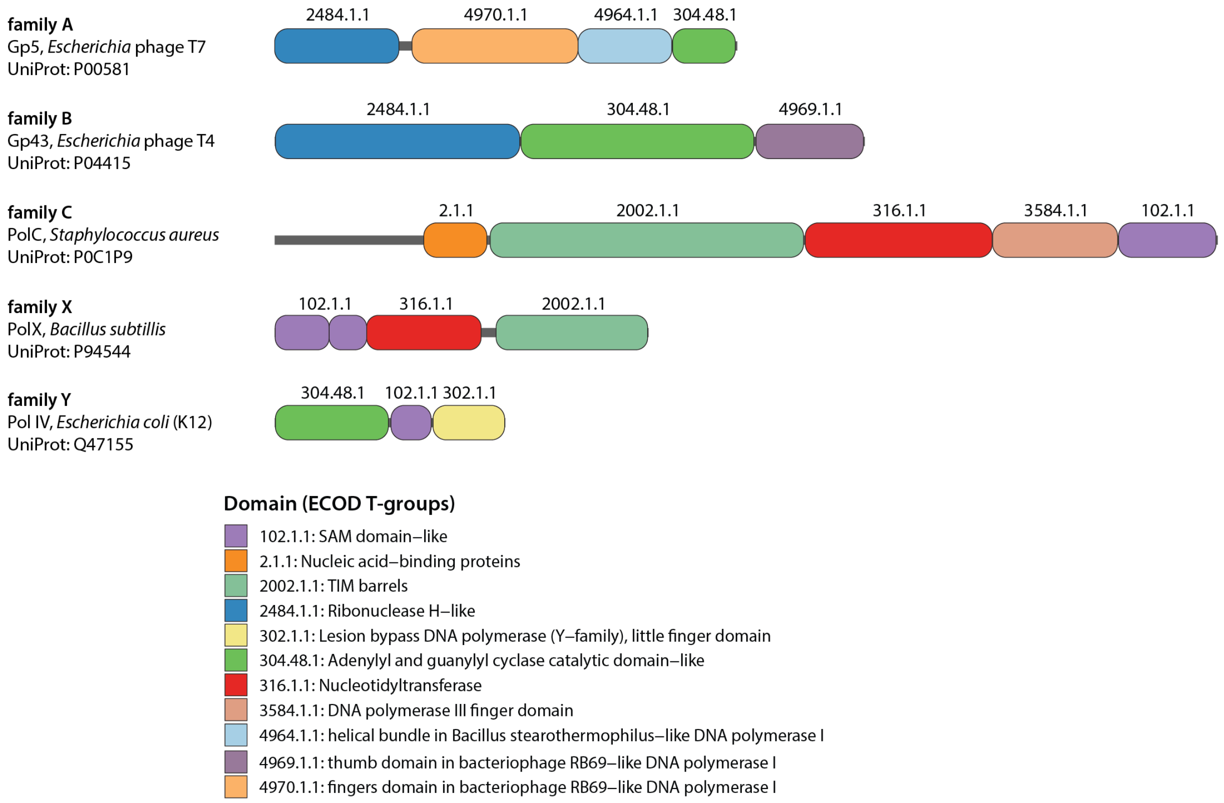


**Figure S6. Domain architectures of representative members of five DNA polymerase families: A, B, C, X and Y.** Domain architectures were obtained with `HHpred` using ECOD as the database. The UniProt accessions of each protein are provided on the left-hand side. Number IDs correspond to ECOD T-groups and colours have been chosen to match those in Figure 5.

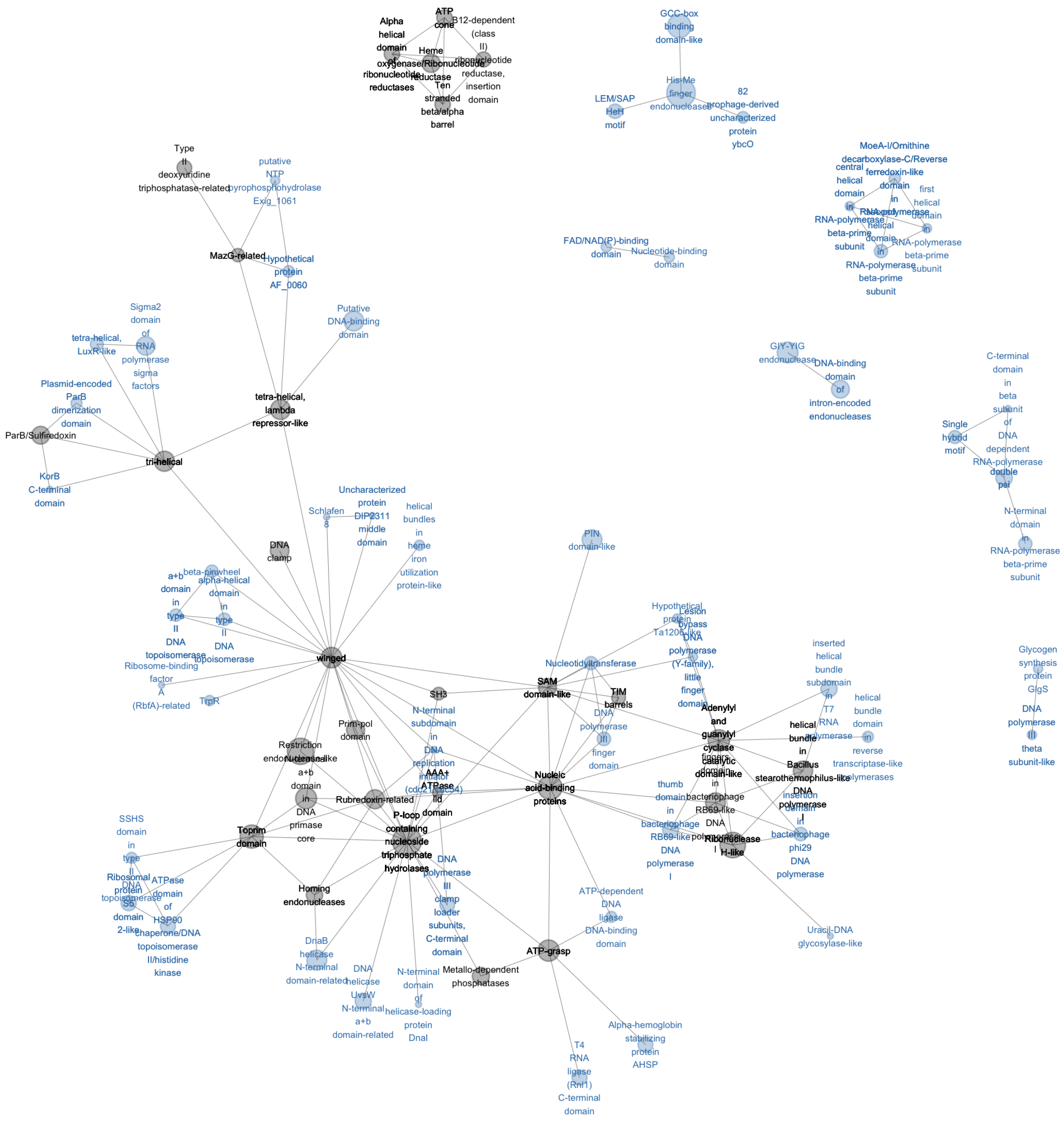


**Figure S7. Domain network of proteins in the 'DNA, RNA and nucleotide metabolism' category.** Nodes in the network represent domains (ECOD T-groups) and are linked by an edge if they co-occur in one rHMM. Size of the node represents its frequency (i.e., number of rHMMs in which it occurs). Colour of the node represents whether the domain occurs in rHMMs assigned to a single functional class (blue) or in rHMMs assigned to multiple functional classes (grey).

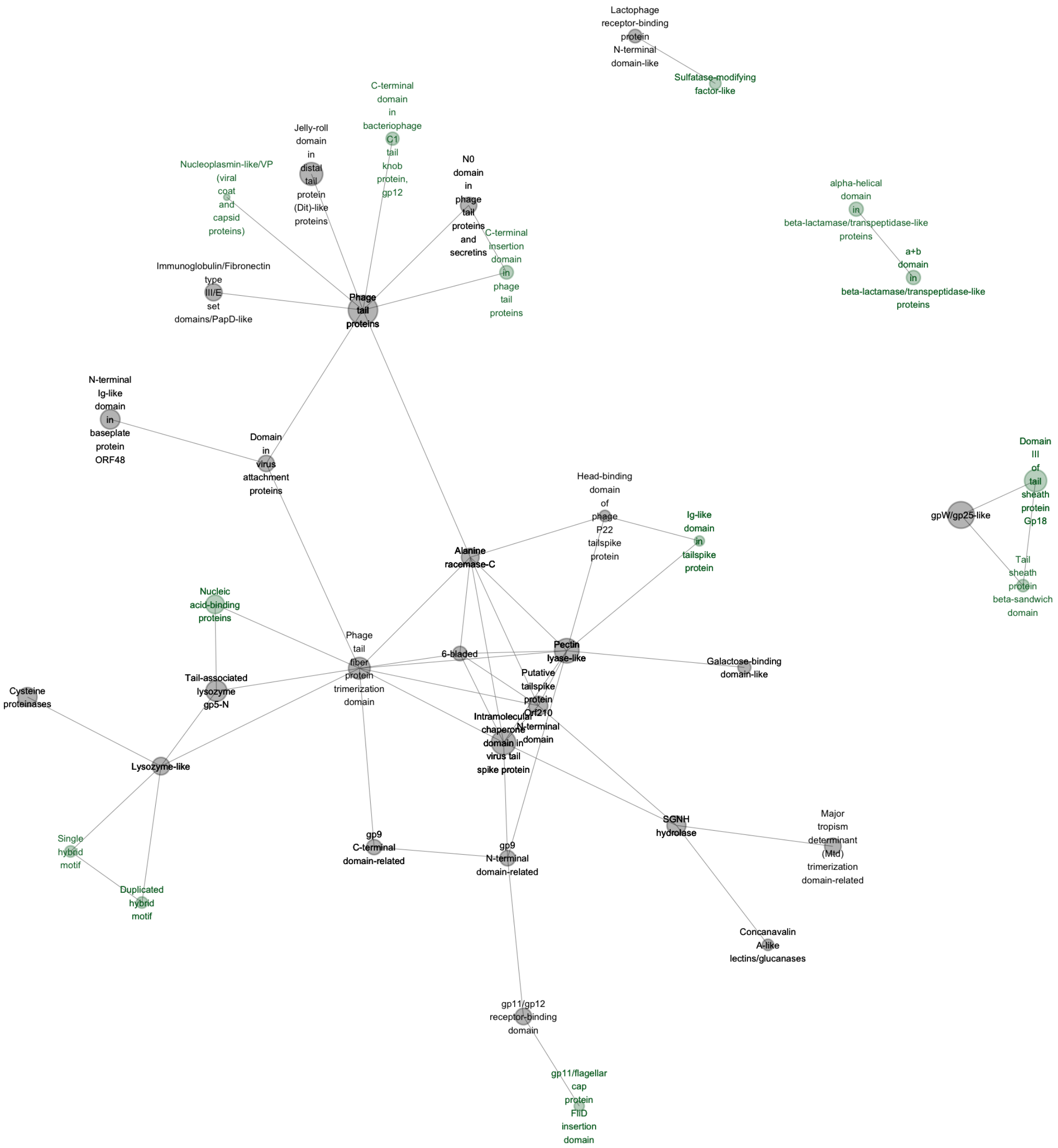


**Figure S8. Domain network of proteins in the 'tail' category.** The network is analogous to the one in Figure S7 with the difference that domains which occur in rHMMs assigned to a single functional class are green instead of blue.

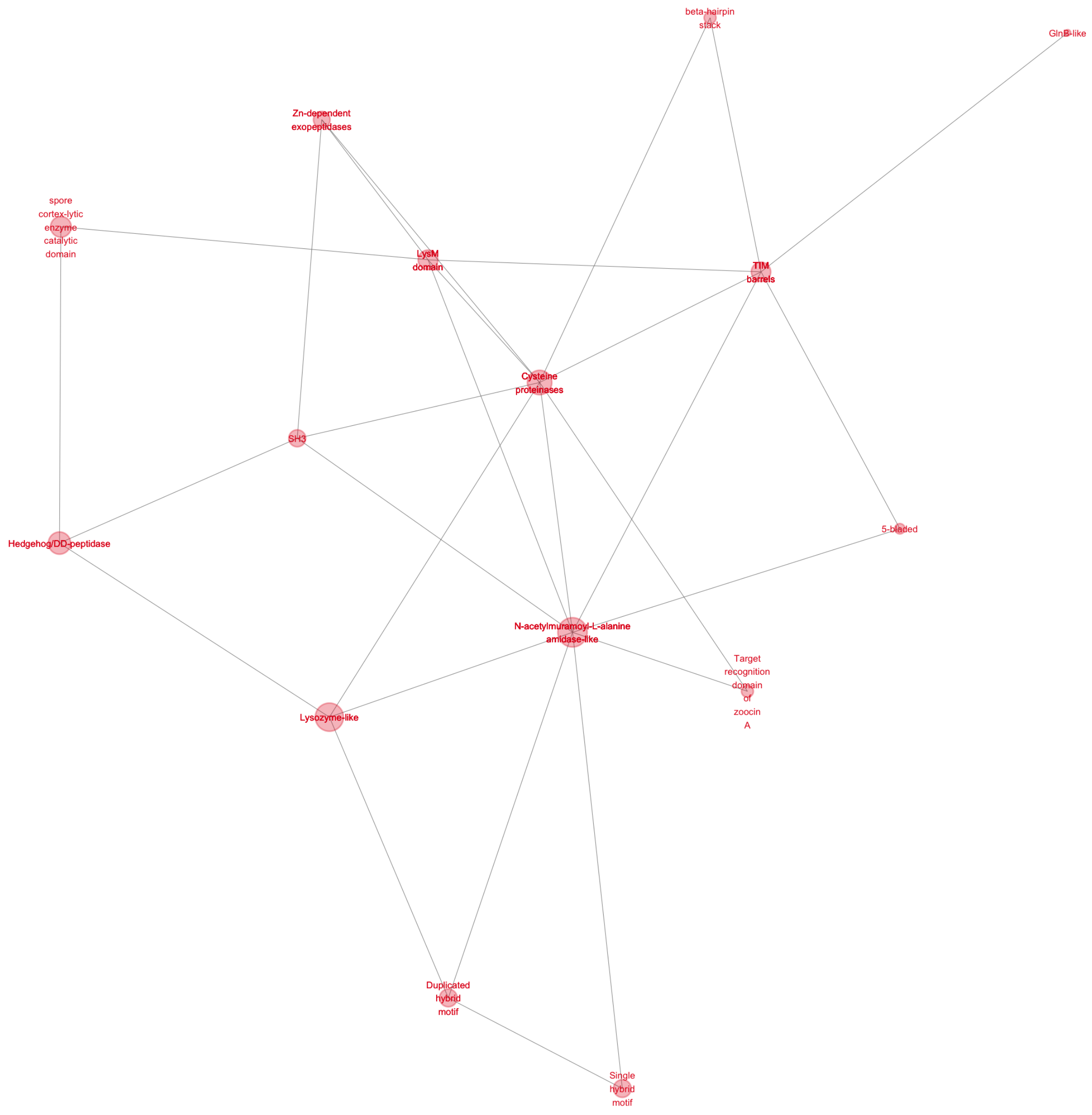


**Figure S9. Domain network of proteins in the 'lysis' category.** The network is analogous to the one in Figure S7 with the difference that domains which occur in rHMMs assigned to a single functional class are red instead of blue.

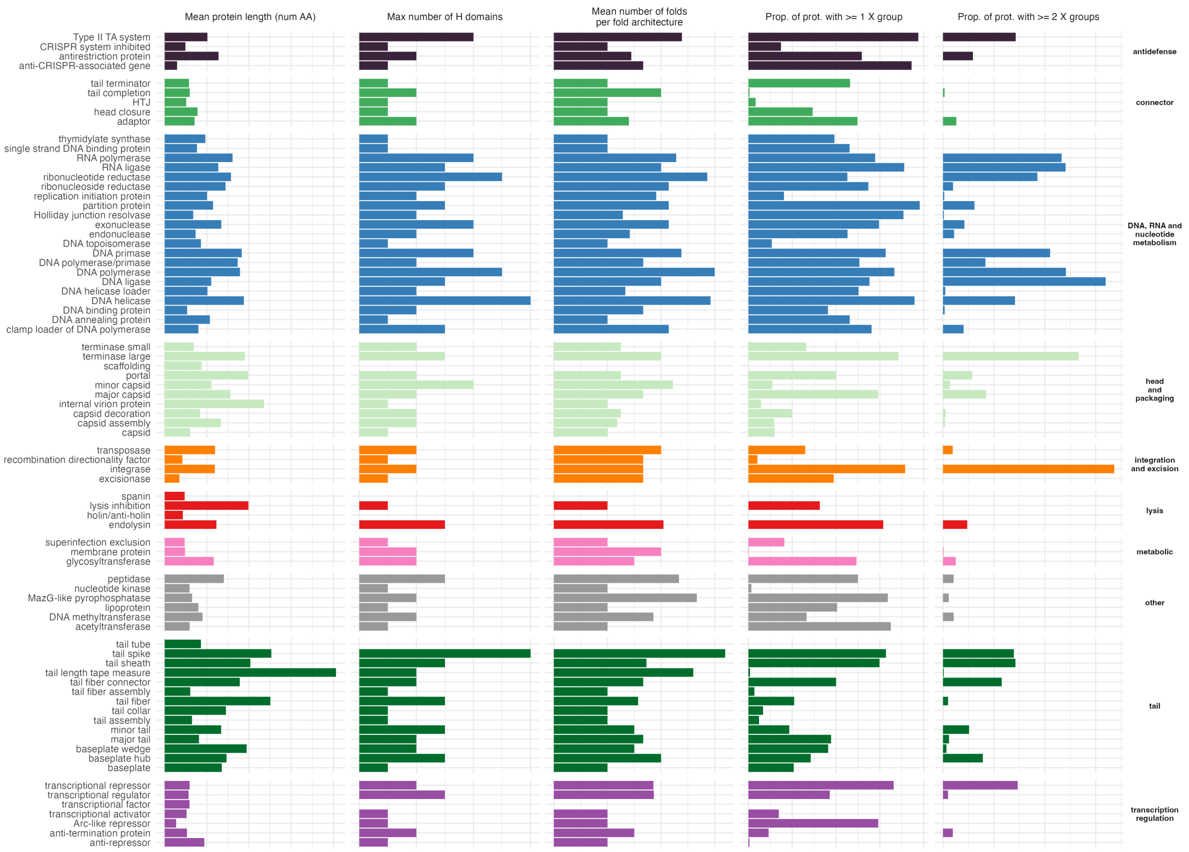


**Figure S10. ECOD coverage per functional class.** Left to right: (i) mean protein length, (ii) maximum number of different domains (H-groups) detected with a single rHMM, (iii) mean number of folds (T-groups) per fold combination, (iv) proportion of proteins with at least one ECOD domain detected at ⩾95% probability and ⩾70% subject coverage, and (iv) proportion of proteins with ECOD domains from at least two distinct X groups, detected at ⩾95% probability and ⩾70% subject coverage. Only the most diverse functional classes (those with ⩾6 rHMM families) are shown.

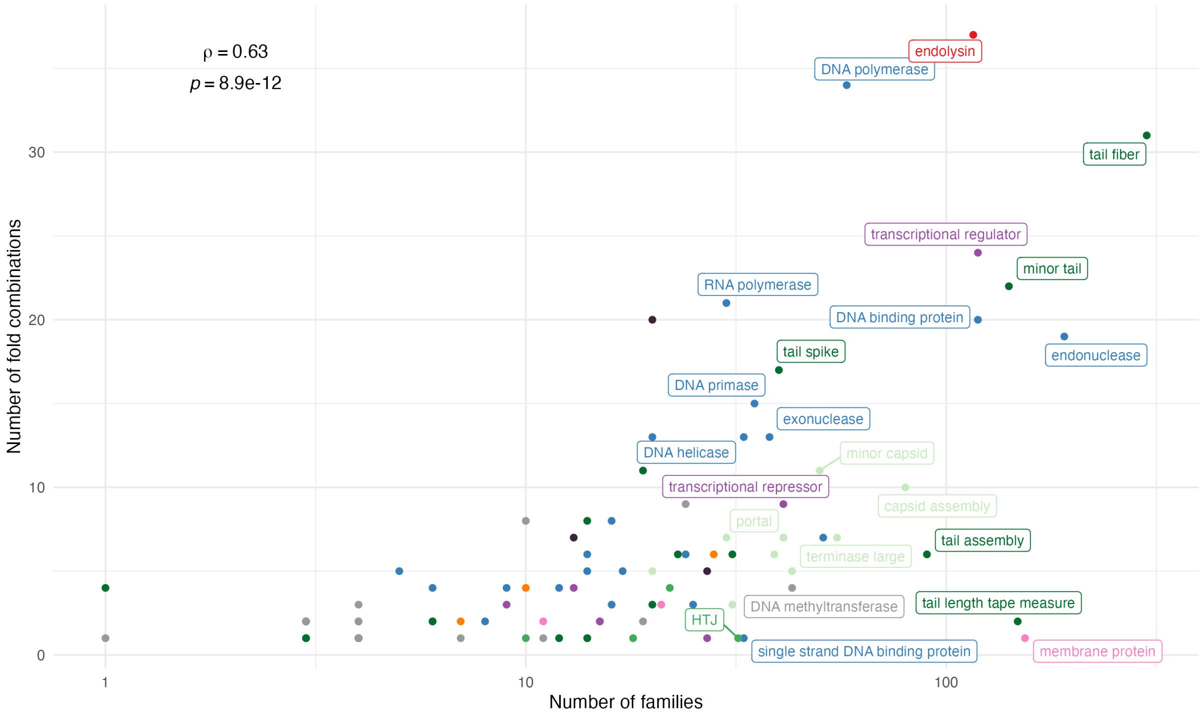


**Figure S11. Fold diversity vs genetic diversity.** Diversity of all functional classes as measured by the number of rHMMs families (X-axis) and the number of fold architectures (i.e., unique combination of ECOD T-groups; Y-axis). Colours correspond to different PHROG functional categories (same as in Figure 2 and Figure 4).

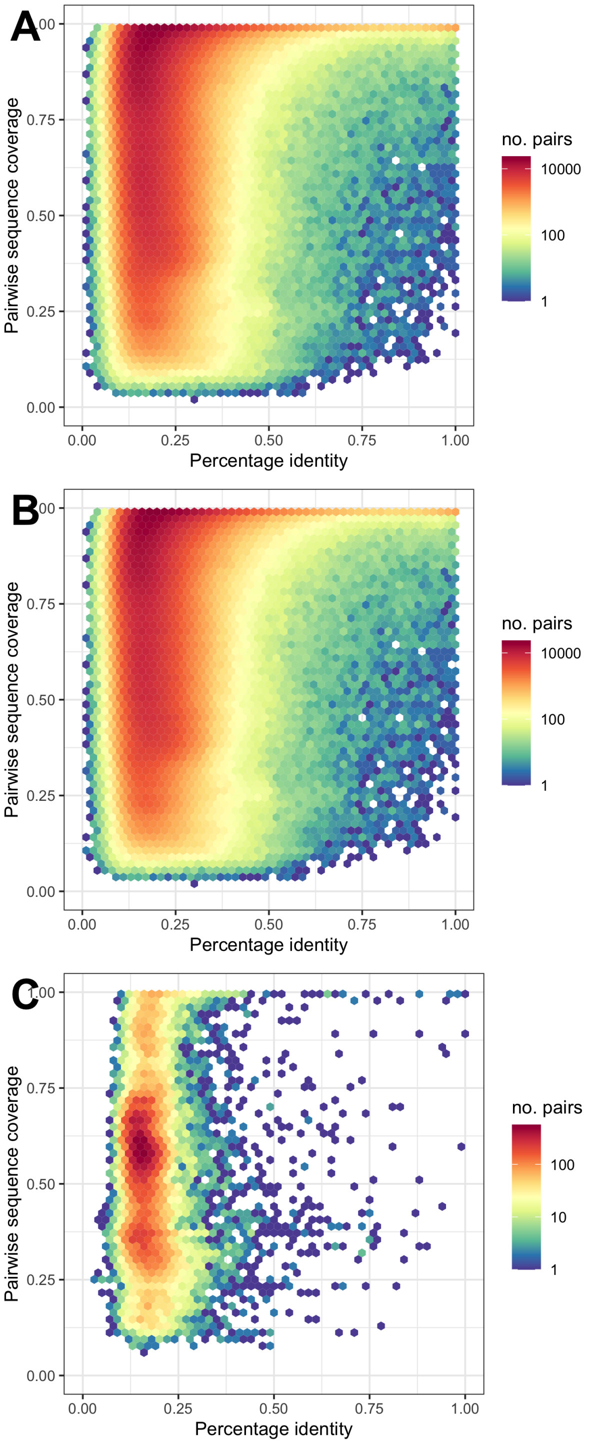


**Figure S12. All-by-all comparison of HMM profiles of rHMMs.** (A) Pairwise sequence coverage (Y-axis) of 133k rHMMs (over $10^{10}$ HMM-HMM comparisons) as a function of pairwise percentage identity. Coverage was calculated as a total coverage based on all hits between a given rHMM pair using hit probability threshold of 95%, and a maximum of query and sequence coverage. Only pairs that overlap on a fragment of $\geqslant$ 50aa are shown. Colour corresponds to the number of rHMM pairs with a given set of parameters, (B) Same as A but with excluded pairs of rHMMs that are detected as mosaic using the domain composition approach, (C) Same as (A) and (B) but only for pairs of rHMMs that are detected as mosaic using the domain composition approach. Note that the vast majority of such pairs shares only very remote homology (no pairwise hit) and is not included in this plot.

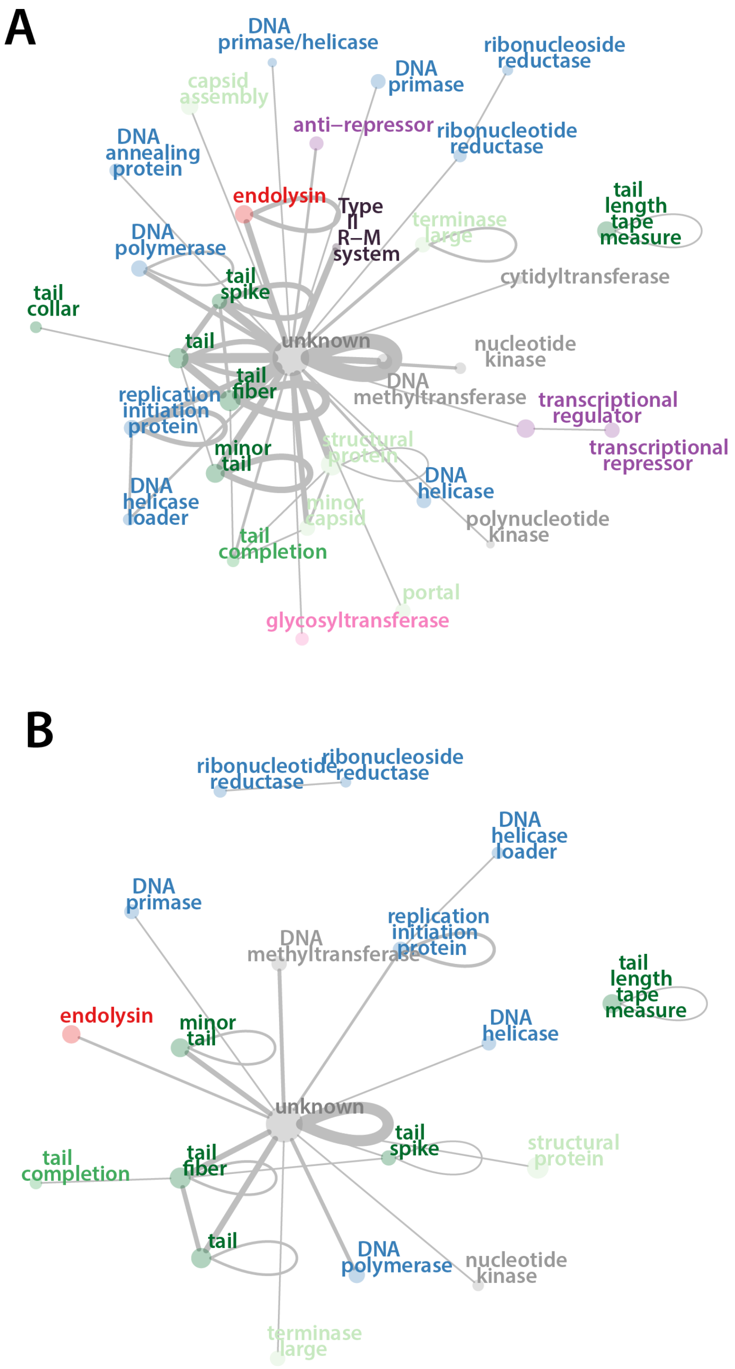


**Figure S13. Networks of functional classes with recent mosaicism.** Nodes represent various PHROG functional classes and the edges connect classes between which there is at least one rHMM pair with evidence of recently emerged mosaicism. Such recently emerged mosaicism between an rHMM pair is defined when two rHMMs show no signal of homology over the majority of their length (at least 50% pairwise coverage) but a strong similarity over a fragment shorter than 50% of their length. By 'strong similarity' we mean percentage identity >= 70% (A) or >= 90% (B). Colour denotes a PHROG category.

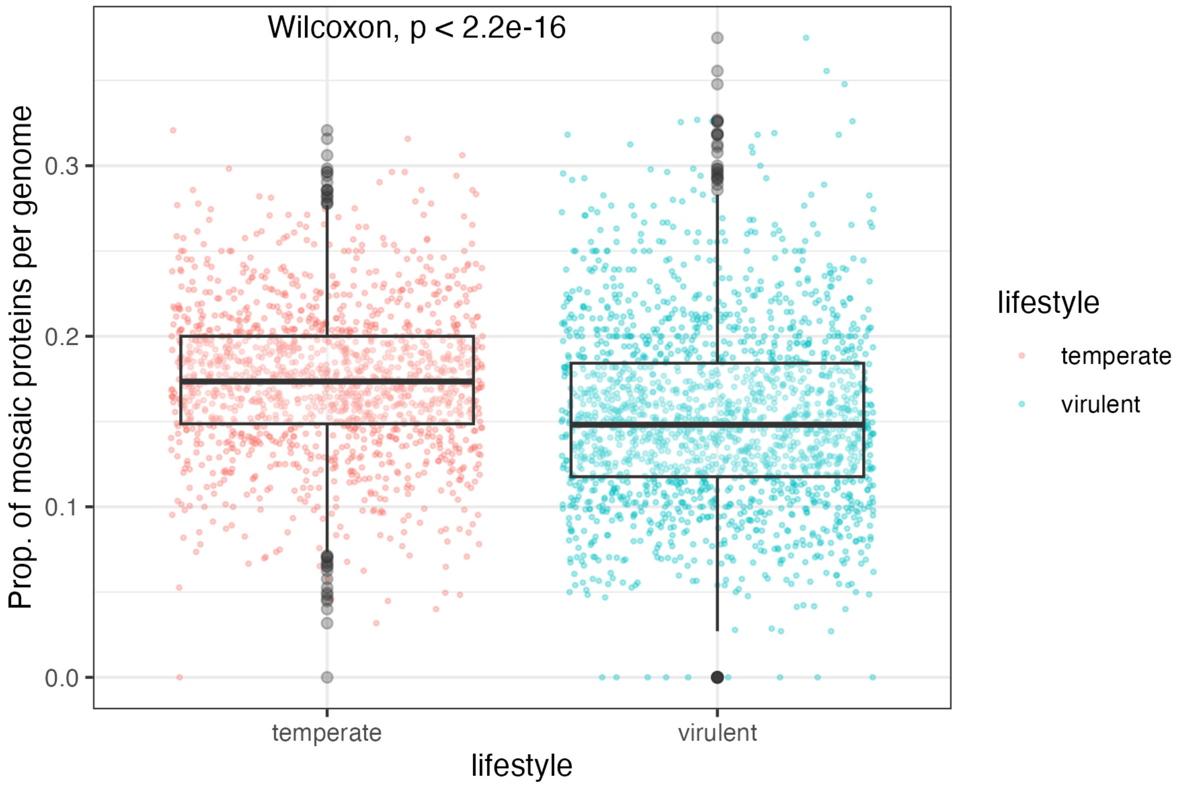


**Figure S14. Proportion of mosaic proteins per genome within temperate and virulent phages.** A protein was defined as mosaic if it belonged to an rHMM that was defined mosaic by ECOD-based or sequence-based definition. The center line within boxplots corresponds to median, lower and upper hinges correspond to the first and third quartiles (the 25th and 75th percentiles). The upper whisker extends from the hinge to the largest value no further than 1.5 * IQR from the hinge (where IQR is the inter-quartile range, or distance between the first and third quartiles). The lower whisker extends from the hinge to the smallest value at most 1.5 * IQR of the hinge. The difference between the groups is statistically significant according to one-sided Wilcoxon rank sum test (W = 963570, $p = 2.8 \times 10^{-45}$).

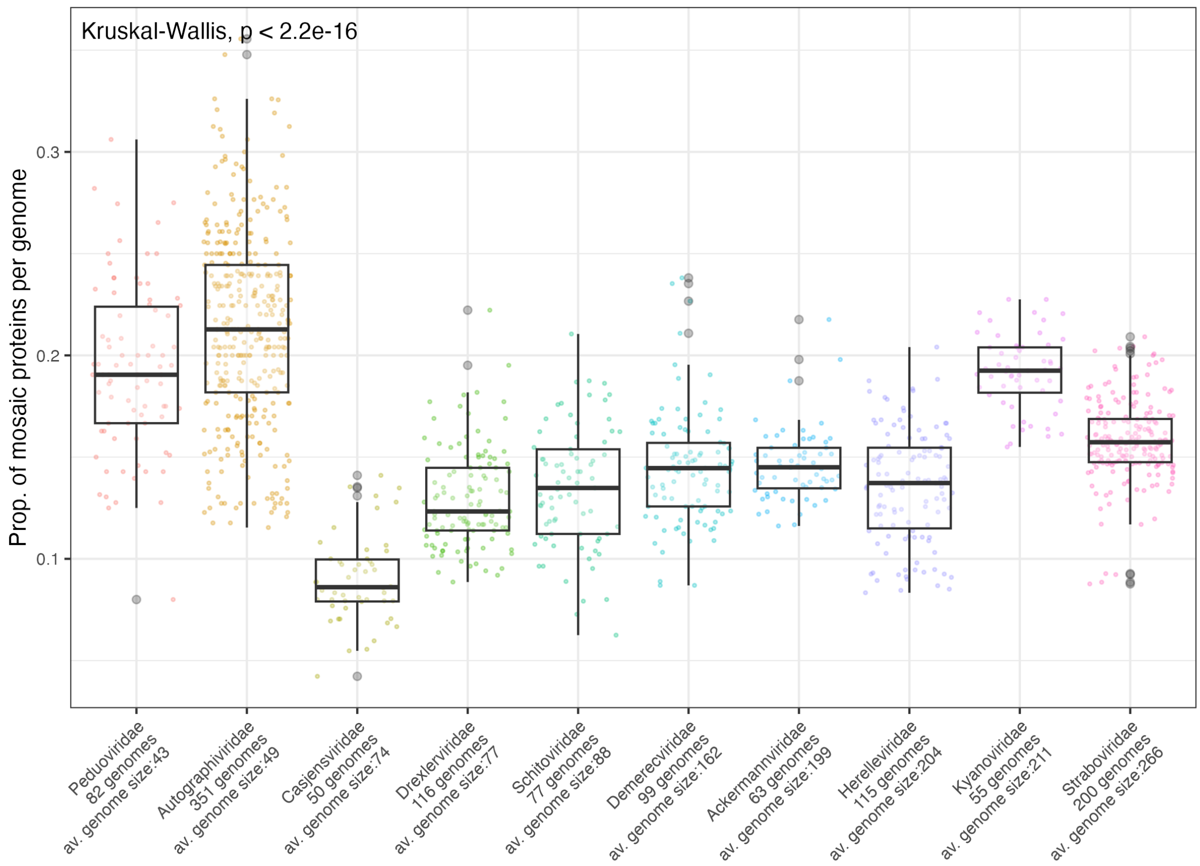


**Figure S15. Proportion of mosaic proteins per genome within phages belonging to different taxonomic families.** The difference between the groups is statistically significant according to Kruskal-Wallis test (Kruskal-Wallis chi-squared = 926.51, df = 40, $p = 2.5 \times 10^{-168}$.). Average genomes sizes represent the median number of genes predicted within the family (rounded up a nearest integer). 'Mosaic protein' is defined the same as in Figure S14. Boxplots are plotted in the same way as in Figure S14.

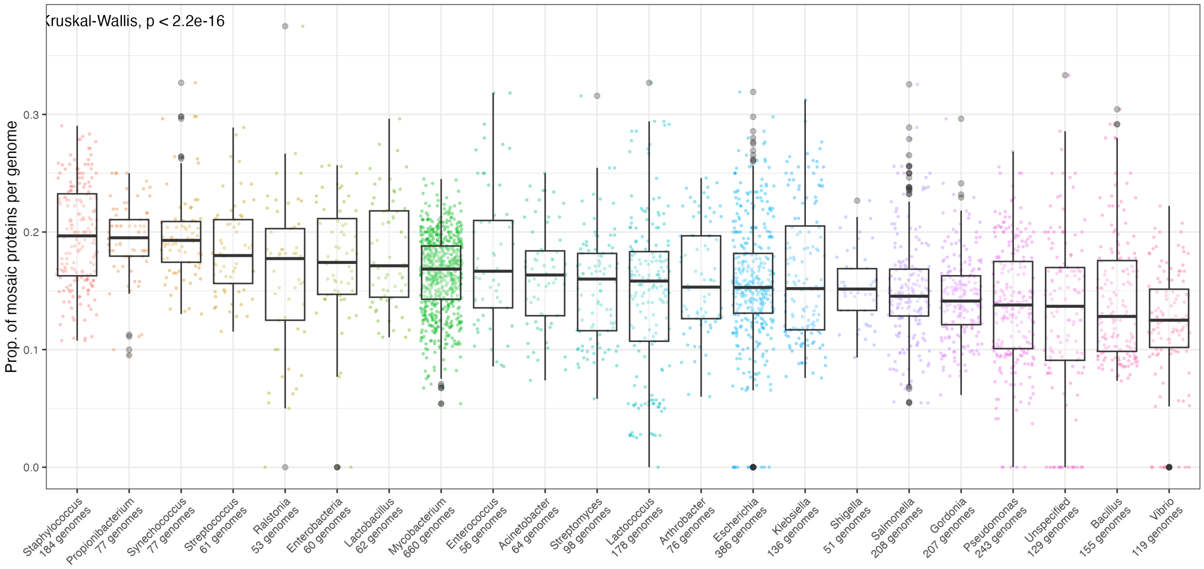


**Figure S16. Proportion of mosaic proteins per genome within phages belonging to different bacterial hosts.** The difference between the groups is statistically significant according to Kruskal-Wallis test (Kruskal-Wallis chi-squared = 878.95, $p = 6.4 \times 10^{-99}$). Average genomes sizes represent the median number of genes predicted within the family (rounded up a nearest integer). 'Mosaic protein' is defined the same as in Figure S14. Boxplots are plotted in the same way as in Figure S14.

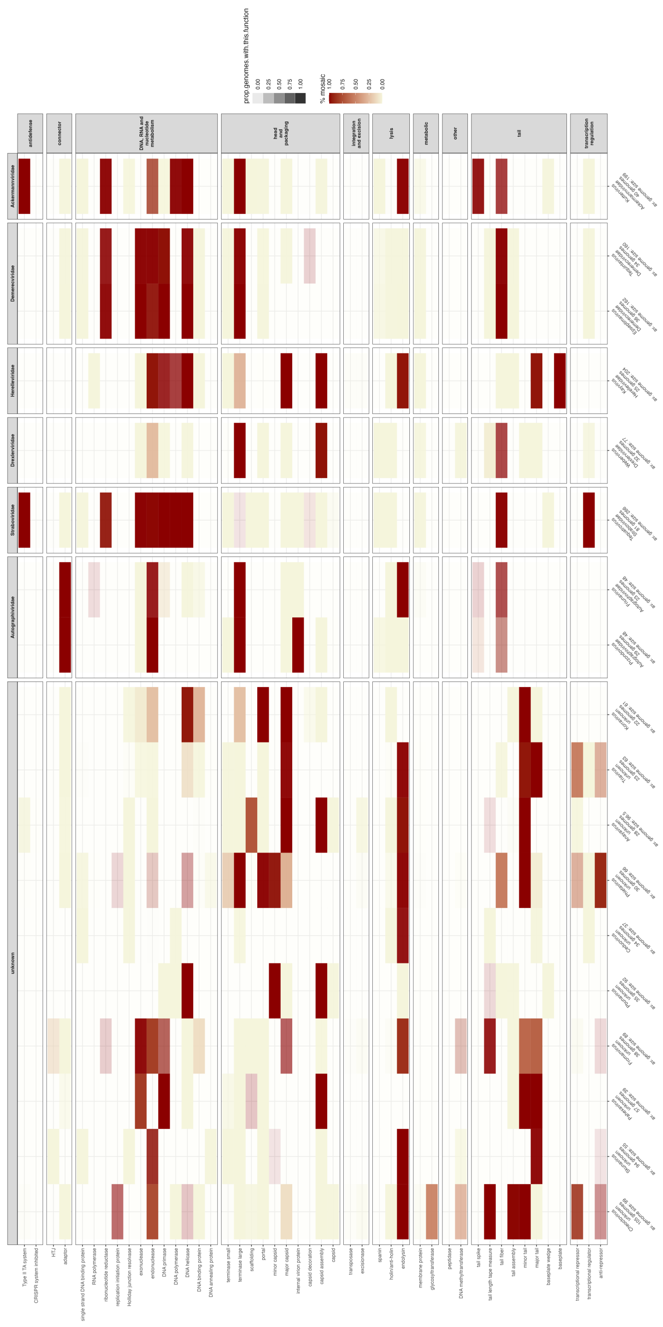

**Figure S17. Mosaicism across taxa.** Mosaicism within multiple protein classes (X-axis) divided into phages of different genera and taxonomic families (Y-axis). Phage genera are grouped into families, and for each genus we show the number of genomes and the average size of these genomes (i.e., the median number of proteins predicted per genome, rounded up to the nearest integer). Transparency denotes the proportion of genomes that include a protein within any rHMM of a given function (note that the function assignment is done very conservatively and many functions will be under-detected). Out of these genomes, we calculated the proportion of those whose protein of a given function was mosaic. This information is represented by the colour (red: very frequently mosaic, beige: rarely mosaic). A protein was defined mosaic if it belonged to an rHMM that was defined mosaic by either domain or sequence-based definition. Only functional classes with genetic diversity of at least 20 families are shown.

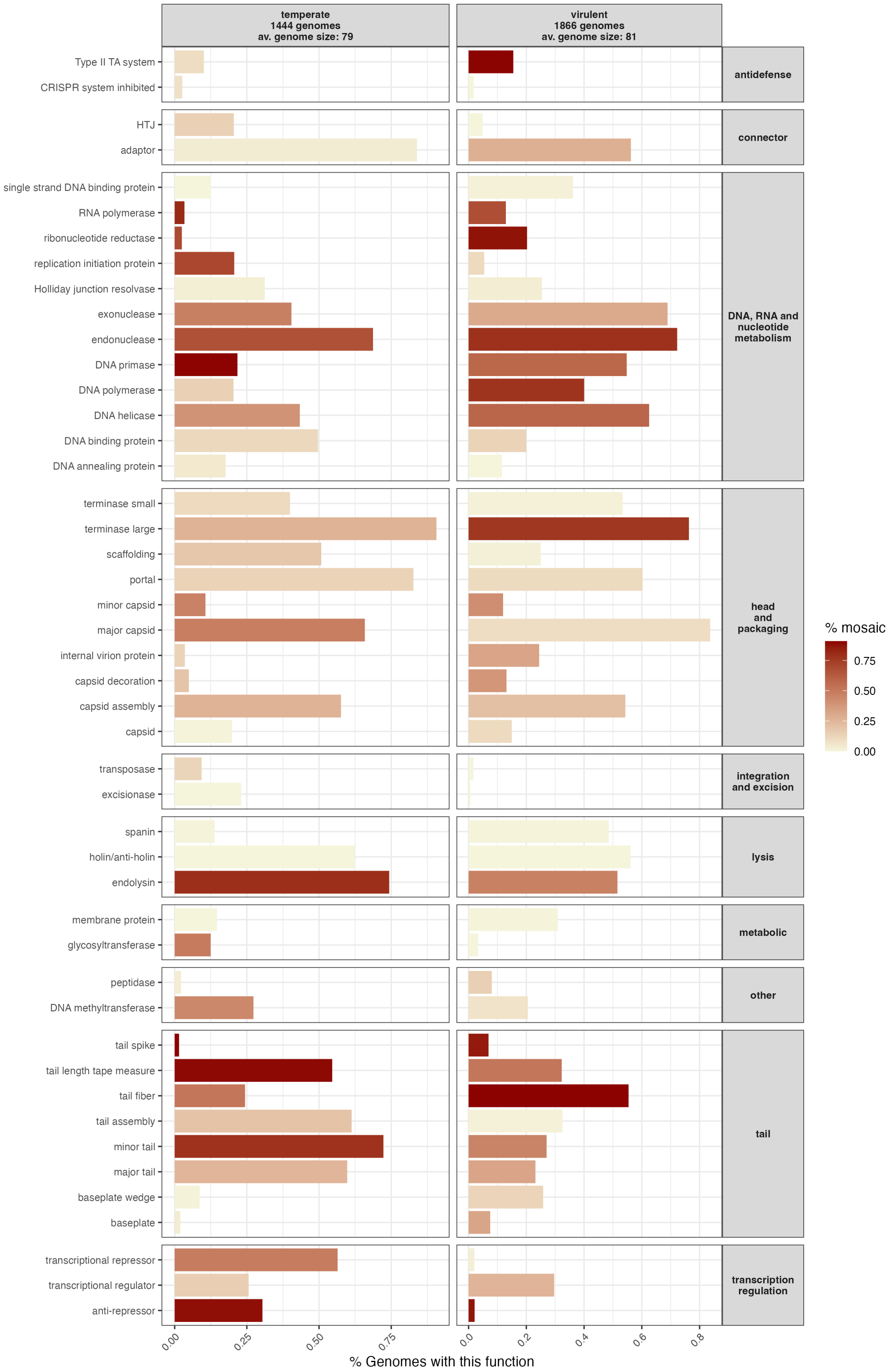


**Figure S18. Mosaicism across lifesyles.** Mosaicism within multiple protein classes (Y-axis) divided into temperate and virulent phages (X-axis, with 1866 virulent and 1444 temperate phages). The length of each bar denotes the proportion of genomes that include a protein within any rHMM of a given function (note that the function assignment is done very conservatively, and many functions will be under-detected). Out of these genomes, we calculated the proportion of those whose protein of a given function was mosaic. This information is represented by the colour (red: very frequently mosaic, beige: rarely mosaic). Only functional classes with genetic diversity of at least 20 families are shown.

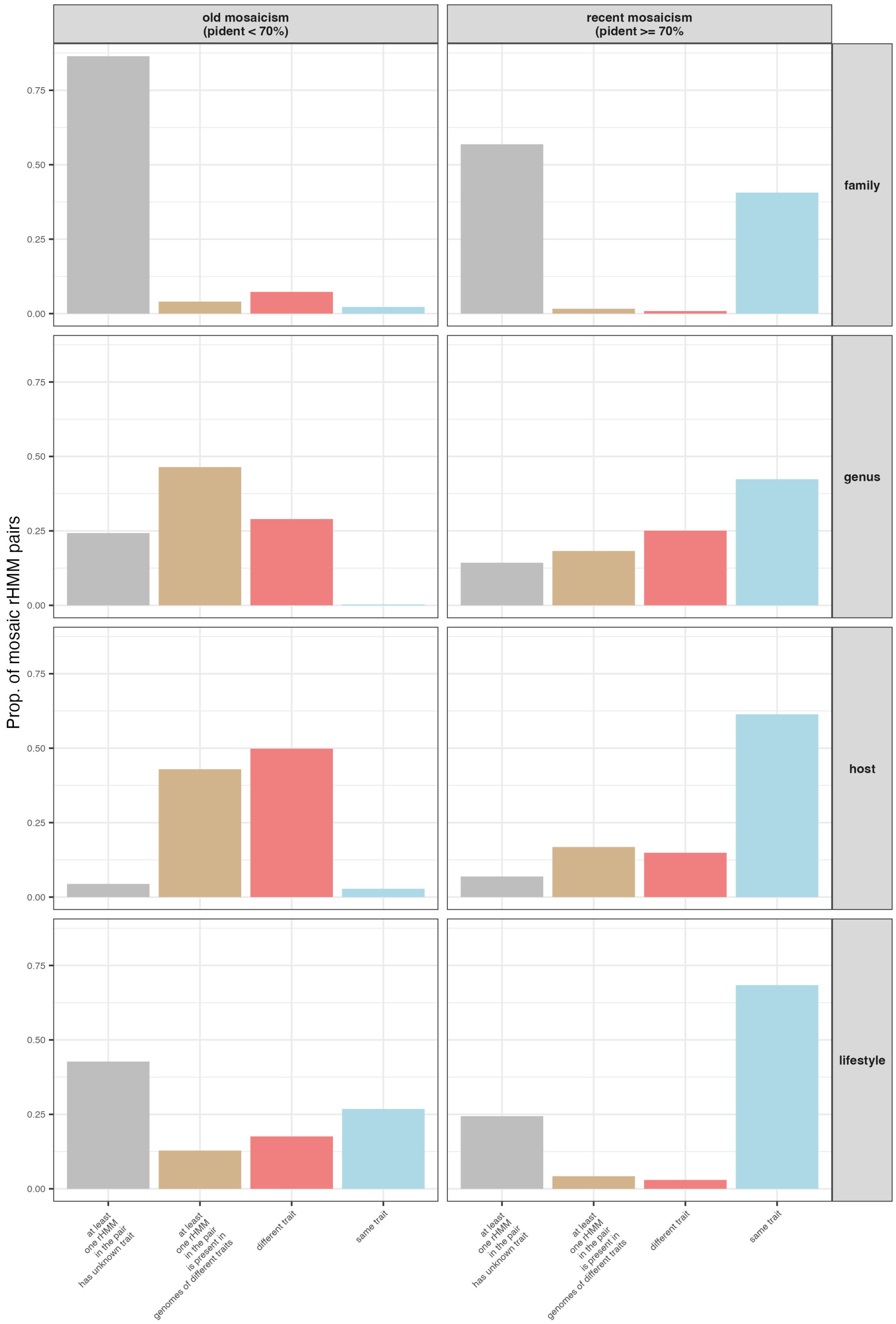


**Figure S19. Recent mosaicism between groups.** Number of mosaic pairs of rHMMs subdivided into old and recently emerged mosaicism (X-axis) that are associated with the same or different trait: taxonomic family, genus, host and lifestyle (Y-axis). A pair of rHMMs was defined mosaic if it was mosaic by either ECOD-based or sequence-based definition. Note that rHMMs may be composed of proteins coming from phages of different groups, e.g. there might be very similar proteins in temperate and virulent phages that are grouped in one rHMM. Such cases are coloured in brown. Additionally, rHMMs composed only of proteins of unknown group (e.g., unclassified taxonomic family) are coloured in grey.

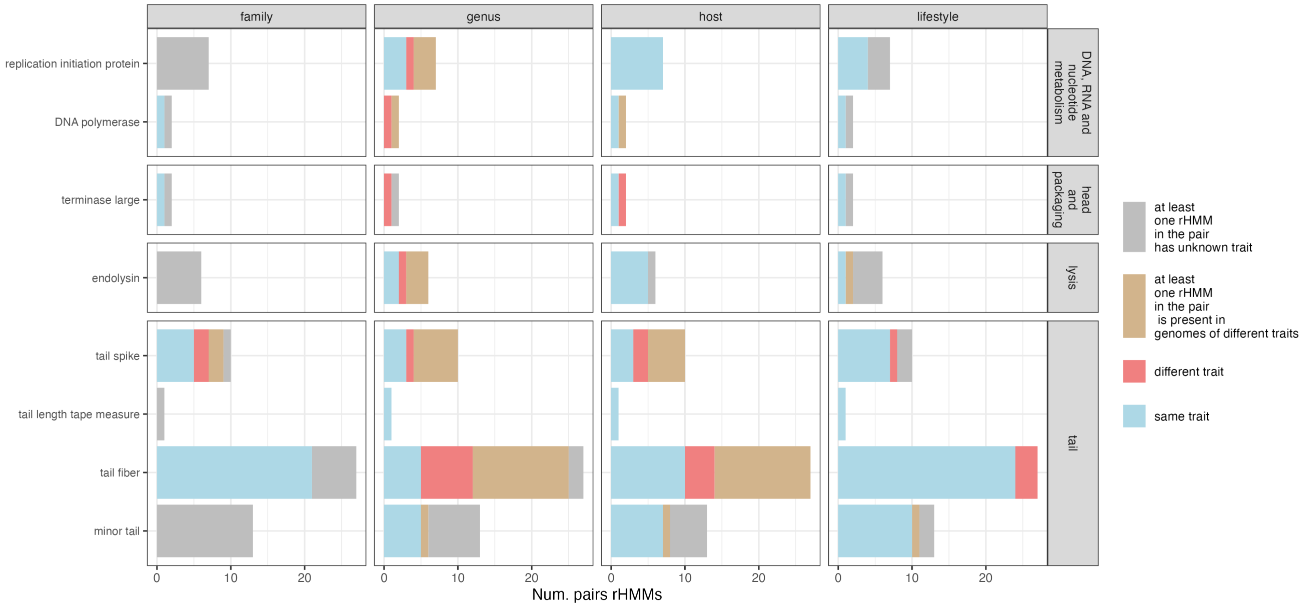


**Figure S20. Recent mosaicism vs. functions.** Number of rHMM mosaic pairs within various functional classes (Y-axis) that has emerged recently (sequence-based mosaicism and pident ⩾70%) and that are associated with the same or different trait: taxonomic family, genus, host and lifestyle (X-axis). Only functional classes with genetic diversity of at least 20 families are shown. Colour legend is the same as in Figure S19.
